# The AMsh glia of *C. elegans* modulates the duration of touch-induced escape responses

**DOI:** 10.1101/2023.12.13.571291

**Authors:** Temitope Awe, Aalimah Akinosho, Shifat Niha, Laura Kelly, Jessica Adams, Wolfgang Stein, Andrés Vidal-Gadea

## Abstract

Once considered mere structural support cells in the nervous system, glia have recently been demonstrated to play pivotal roles in sensorimotor processing and to directly respond to sensory stimuli. However, their response properties and contributions to sensory-induced behaviors remain little understood. In *Caenorhabditis elegans*, the amphid sheath glia (AMsh) directly respond to aversive odorants and mechanical stimuli, but their precise transduction machinery and their behavioral relevance remain unclear.

We investigated the role of AMsh in mechanosensation and their impact on escape behaviors in *C. elegans*. We found that nose touch stimuli in immobilized animals induced a slow calcium wave in AMsh, which coincided with the termination of escape reversal behaviors. Genetic ablation of AMsh resulted in prolonged reversal durations in response to nose touch, but not to harsh anterior touch, highlighting the specificity of AMsh’s role in distinct escape behaviors.

Mechanotransduction in AMsh requires the α-tubulin MEC-12 and the ion channels ITR-1 and OSM-9, indicating a unique mechanosensory pathway that is distinct from the neighboring ASH neurons. We find that GABAergic signaling mediated by the GABA-A receptor orthologs LGC-37/8 and UNC-49 play a crucial role in modulating the duration of nose touch-induced reversals.

We conclude that in addition to aversive odorant detection, AMsh mediate mechanosensation, play a pivotal role in terminating escape responses to nose touch, and provide a mechanism to maintain high sensitivity to polymodal sensory stimuli.

**Significance:** Polymodal nociceptive sensory neurons have the challenge of multitasking across sensory modalities. They must respond to dangerous stimuli of one modality, but also adapt to repeated nonthreatening stimuli without compromising sensitivity to harmful stimuli from different modalities. Here we show that a pair of glia in the nematode *C. elegans* modulate the duration of nose-touch induced escape responses. We identify several molecules involved in the transduction of mechanical stimuli in these cells and show that they use the signaling molecule GABA to modulate neural function. We propose a mechanism through which these glia might function to maintain this polysensory neuron responsive to dangerous stimuli across different modalities.

## Introduction

To survive and thrive animals must quickly respond to dangerous stimuli in their environment. Escape circuits are often streamlined and centralized so that dangerous stimuli across multiple sensory modalities can access and trigger a potentially life-saving escape response with minimal delay. For example, either visual or mechanical stimulation of the head of a crayfish can cause the medial giant neurons to trigger an escape response (Edwards *et al*., 1999). In amphibians and fish, the Mauthner neurons can be similarly stimulated to produce an escape response in response to visual, chemical, or mechanical stimuli (Hackett et al., 1986). In the nematode *C. elegans*, the ASH neurons respond to noxious chemical, osmotic, and mechanical stimuli, initiating an escape response when activated by one of these stimuli (de Bono and Maricq, 2005). One challenge presented by convergent multimodal circuits such as those described above is how to adapt to repeated stimuli from one modality without compromising sensitivity to harmful stimuli from another. Recent work on the nematode *C. elegans* has revealed that glia may play a role in addressing this challenge.

Glia are essential non-neuronal components of most nervous systems. They regulate numerous aspects of neural function including neurogenesis, neuronal migrations, axonal growth, synapse formation, and circuit function (Kuwada, 1986, Pomeroy and Purves, 1988, Song *et al*. 2002, Merkle *et al*. 2004 Funfschilling *et al*. 2012, Allen and Lyons, 2018). A surprising finding of the last decade is that glia can contribute to sensory processing (Yadav *et al*. 2019; Duan *et al*. 2020; Fernandez-Abascal *et al*. 2022; Chen *et al*. 2022) and respond directly to external stimuli (Paemeleire and Leybaert 2000, Ding *et al*. 2015, Duan *et al*. 2020). There is also evidence that glia can communicate with, and influence processing in, neurons via paracrine excitatory and inhibitory chemical signaling (Araque *et al*. 2014; McClain *et al*. 2014; Costa and Moreira, 2015; Belzer and Hanani, 2019), although much of these actions remain little understood. In the nematode *Caenorhabditis elegans (C. elegans*), a bilateral pair of amphid sheath glia (AMsh) play an integral role in the sensory processing of the amphid sense organs. One AMsh glia and 12 sensory neurons comprise each of these organs. They are found in the head of the worm (review: Shaham 2016) and mediate chemosensation, thermosensation, osmosensation, and magnetosensation (Driscoll and Tavernarakis, 1997; Bargmann, 2006; Vidal-Gadea *et al*. 2015). Ablation of the AMsh causes developmental impairments in amphid neurons (Heiman and Shaham, 2009; Singhvi *et al*. 2016) and the behaviors associated with these sensory modalities (magnetotaxis: Vidal-Gadea *et al*. 2015, chemotaxis: Bacaj *et al*. 2008, thermotaxis: Yoshida *et al*. 2016, mechanosensation: Fernandez-Abascal *et al*. 2022).

Duan *et al*. (2020) showed that AMsh autonomously respond to aversive odorants via the G-protein-coupled receptor (GPCR) GPA-3. This pathway involves the Inositol Triphosphate Receptor ITR-1 and OSM-9, the *C. elegans* homolog of the mammalian transient receptor potential channels of the vanilloid subtype 4 (TPPV4). AMsh responses appear to mediate the adaptation of avoidance behaviors to repeated aversive odorant exposure by releasing GABA onto the two polymodal ASH neurons. The AMsh and the ASH neurons also respond to mechanical stimulation of the anterior end of the worm (“nose touch”). No other amphid organ neurons are activated by nose touch (Kaplan and Horvitz, 1993; Hilliard *et al*. 2005; O’Hagan and Charlie 2005; Ding *et al*. 2015), and two lines of evidence indicate that nose touch transduction in AMsh and the ASH neuron occurs independently: First, the AMsh display higher sensitivity to mechanical stimulation than ASH neurons (Ding *et al*. 2015). Second, in the ASH neurons, nose touch is transduced by amiloride-sensitive DEG/ENAC channels (Geffeney *et al*. 2011) and can be blocked with the epithelial sodium channel blocker amiloride. In contrast, nose touch responses of the AMsh are amiloride-insensitive (Ding *et al*. 2015; Fernandez-Abascal *et al*. 2022).

The transduction mechanism underlying AMsh nose touch sensitivity remains unresolved, as does the behavioral relevance of the AMsh’s response. The latter is particularly intriguing because nose touch responses spread very slowly in AMsh, taking over one minute from the terminals near the amphid organ to the soma that is a few micrometers away (Ding *et al*. 2015). This delay puts the AMsh response outside of the timeframe of any previously described and ecologically relevant touch-induced *C. elegans* behavior.

To understand the role of the AMsh in mechanosensation and adaptive behavior, we recorded nose touch-induced AMsh activation in restrained and freely crawling animals. We find that AMsh modulated reversal behaviors elicited by nose touch. Nose touch activated AMsh and the peak of the AMsh activity coincided with the end of nose touch-induced reversals. Reversal duration was modulated through AMsh GABA signaling and the GABA_A_ receptor orthologs LGC-37, LGC-38, and UNC-49. We found that AMsh touch sensitivity relies on the IP3 (ITR-1) and TRPV4 (OSM-9) channels, demonstrating a convergence of mechanisms between odor detection and mechanosensation in AMsh. We hypothesize that AMsh function to maintain the polysensory ASH neurons responsive to dangerous stimuli across multiple sensory modalities.

## Results

### Nose touch elicits a slowly spreading calcium wave in AMsh

To corroborate mechanosensitivity in AMsh, we used GCaMP6s to measure AMsh calcium responses to nose touch in glue-immobilized animals (Fig.1a). Using a glass probe we delivered mechanical nose stimuli averaging 22.5±4.8 μm displacement and lasting 1.2±0.4 seconds (N=13). Consistent with Ding *et al*. (2015), we found that touch stimuli induced a calcium wave in the AMsh glia terminals that propagated slowly toward the soma, taking an average of 76.6±4.9 seconds from the onset of stimulus for the signal to peak in the soma (N=8 of 13 animals, Fig. 1b,c). In 5 animals, nose touch initiated a calcium wave in the terminals, but it failed to propagate to the soma.

**Figure 1.**
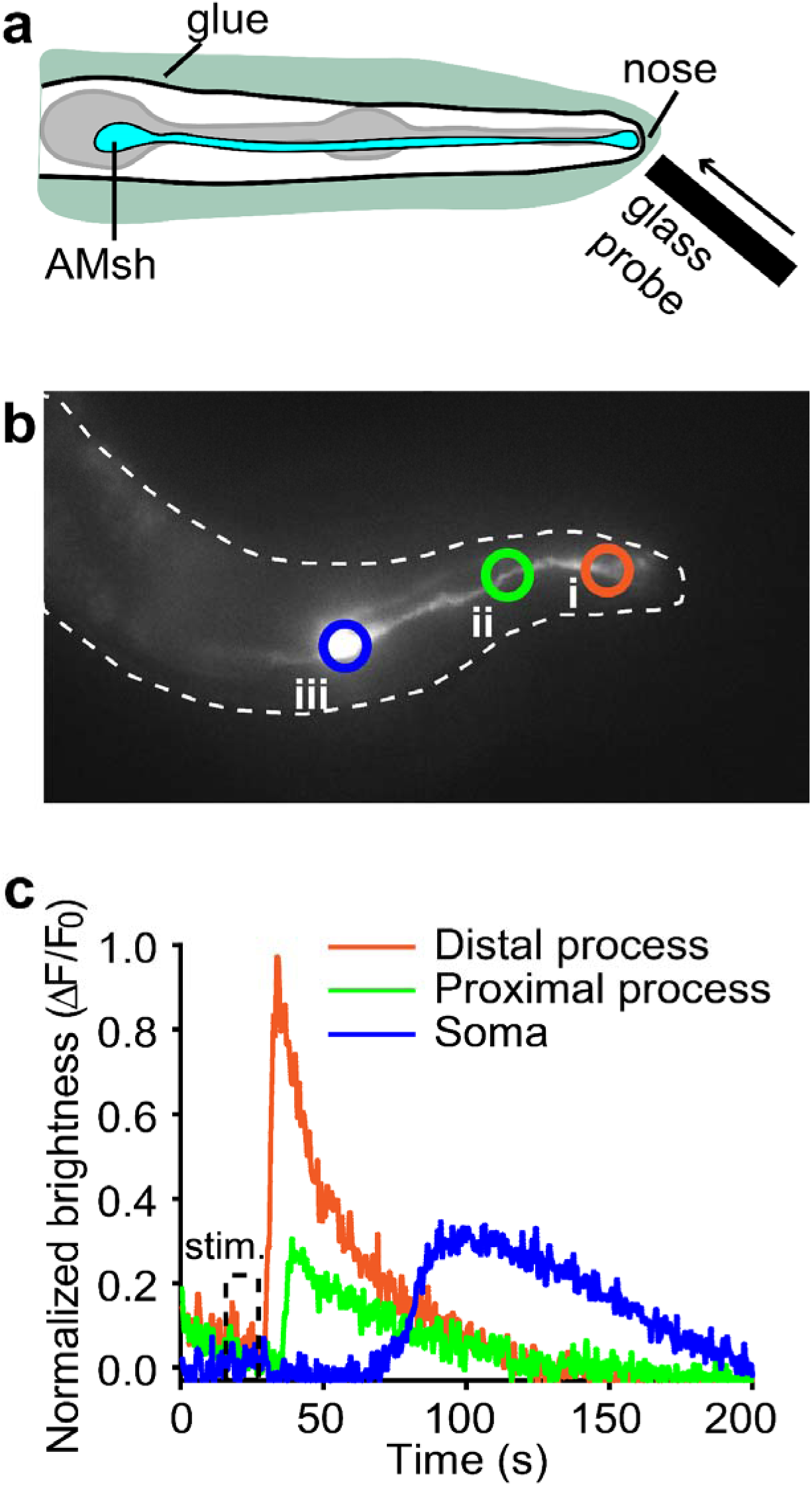
The AMsh glial cells respond with a slow calcium wave to mechanical stimuli in immobilized worms. **a)** Experimental approach. Worms expressing GCaMP6s in AMsh glia were immobil zed on agar pads with WORMGLU (GluStitch) and stimulated with a glass pipette. **b)** Image of the anterior end of a day-1 adult worm expressing GCaMP6s throughout AMsh. The circles show different points along the processes of the glia, starting at the distal end (i) moving along the process all the way to the soma (iii). **c)** Mechanical stimulation (stim.) elicited a rapid increase in calcium fluorescence in the distal process of AMsh. Calcium fluorescence slowly spread towards the soma where it peaked at ∼100 seconds post-stimulation.

### AMsh calcium signals peak at the end of nose touch-induced reversals

To investigate whether AMsh activity is associated with natural behaviors worms perform during locomotion we measured AMsh calcium activity in freely crawling worms in a 30-second assay with minimal mechanosensory stimuli. To correlate AMsh activity with ongoing behaviors, we recorded AMsh soma fluorescence for 30 seconds and then searched for phases of high activity within this time frame. High activity was defined as exceeding the average fluorescence of the 30-second trial by 1 SD. For comparison between animals, all data were normalized to the maximum background fluorescence.

We then classified the animal’s behavior into three different states: i) forward crawling, ii) reversing (backwards crawling), and iii) deep bending (performing deep bend turns). We determined the duration of each behavior and then compared the percentage of time during the behavior when the AMsh calcium activity exceeded the average fluorescence by 1 SD (Fig. S1a). We found no clear difference between behaviors, suggesting that high AMsh activity is not differentially regulated during these behaviors. Similarly, there was no difference between behaviors when we compared the absolute duration of high AMsh fluorescence during each of the aforementioned behaviors (Fig. S1b) and the frequency with which phases of high activity occurred (Fig. S1c).

To test the potential contribution of AMsh activity to behaviors elicited by mechanical stimuli, we applied nose touch and harsh anterior touch (henceforth “harsh touch”) stimuli. Nose touch and harsh touch differ in strength, and location where they are applied. They further differ in that harsh touch is delivered to the animal, while nose touch is the result of the animal colliding with an immobile object placed along its path. They also differ in the sensory pathways activated. Nevertheless, both stimuli elicit a reversal escape behavior in freely moving animals (Kaplan and Horvitz 1993). Nose touch involves the animal initiating contact by crawling against a stationary obstacle. In our experiments, we thus manually placed an obstacle (i.e., a platinum worm pick) in the path of crawling worms to facilitate a collision (Fig. 2a). For harsh touch, we pressed a platinum wire pick against the anterior region of the worm’s body.

**Figure 2.**
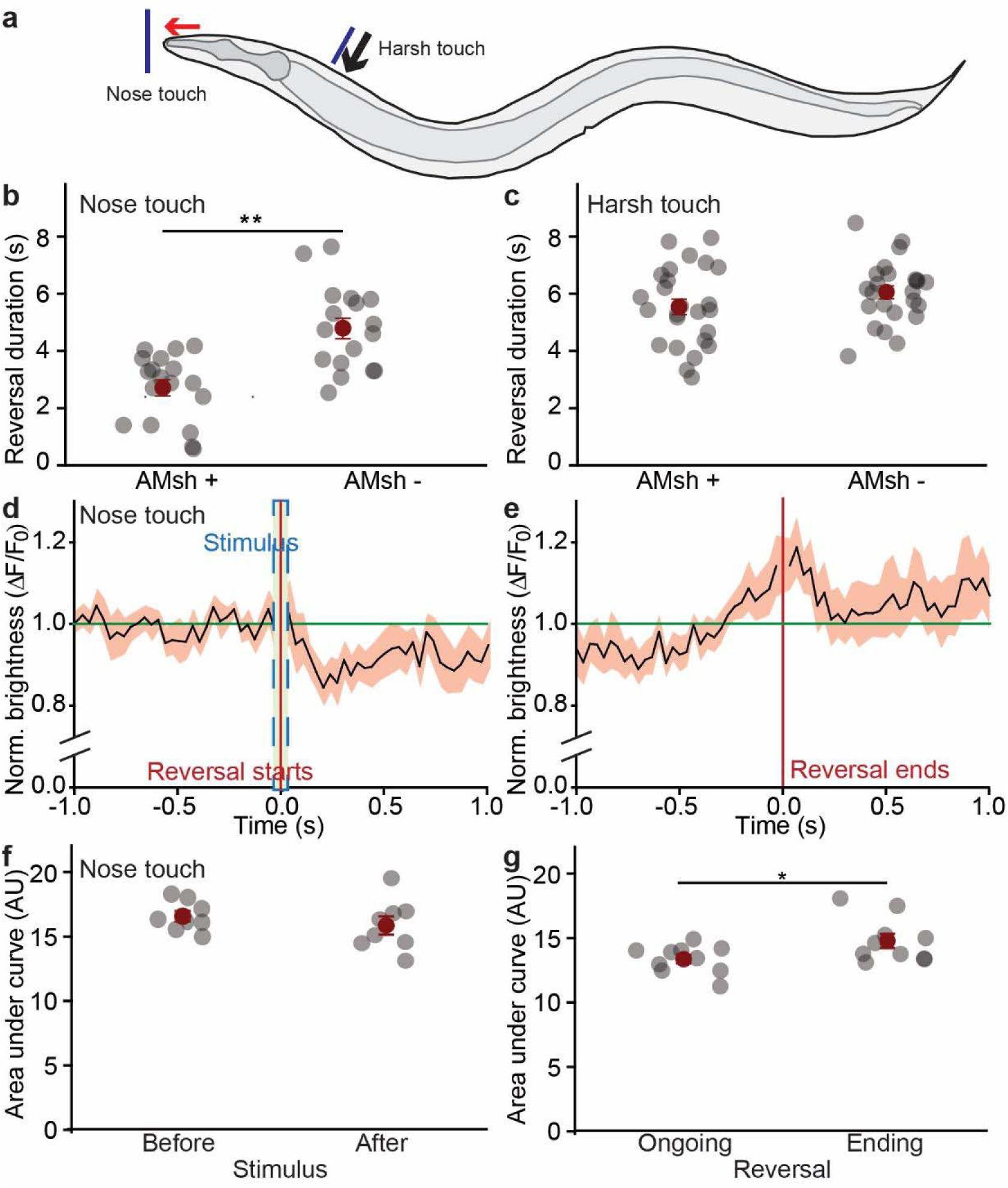
AMsh activity peaks at the end of nose touch in freely crawling worms. **a)** Diagram illustrating two anterior mechanical stimuli used in this study (nose touch and harsh touch). **b)** Reversal duration for AMsh-ablated animals in response to nose touch, and harsh touch **(c).** Data are mean ± SEM. *P < 0.05, **P < 0.001, student’s t-test. Sample sizes are N≥17. **d)** AMsh brightness over time (normalized to background brightness). GCaMP6s brightness in AMsh is shown before and after the administration of mechanical stimulations to the nose (occurring during the period within the blue dashed box). Worms initiated a rapid escape reversal whose onset is marked by the red vertical line. **e)** MaxiAMsh brightness before and after the end of reversals triggered by a nose touch stimuli. The red vertical line denotes the moment worms ceased backward motion and resumed forward motion. **f)** Comparison of the area under the curve for AMsh GCaMP6s brightness for the 0.5 seconds immediately before and immediately after the delivery of nose touch stimuli. **g)** Comparison of the area under the curve for AMsh GCaMP6s brightness during the 1 second spanning the end of the reversal (marked by the vertical red line), and the 0.5 seconds preceding and following this time. Data are mean ± SEM. *P < 0.05, paired student’s t-tests. N=11.

We recorded and measured the duration of the induced reversals and then compared freely crawling wild-type worms with worms where the AMsh had been ablated genetically (through the targeted expression of a caspase, see methods). No difference was seen in response to anterior harsh touch (controls: 5.8±0.3 s; ablated: 6.3±0.2 s, t-test, p=0.16, Fig. 2d). However, AMsh-ablated worms displayed significantly longer reversal durations in response to nose touch (control: 2.7±0.3 s; ablated: 4.8±0.4 s, t-test, p<0.001, Fig. 2b), suggesting a potential contribution of AMsh activity to this specific behavior.

We also recorded GCaMP6s fluorescence from the AMsh cell body of freely crawling worms in response to nose touch and tracked it throughout the execution of the escape reversal. As expected, nose touch did not result in immediate AMsh soma activation, and fluorescence remained initially unaltered (Figs. 2d, f). However, AMsh activity peaked the moment backward locomotion ended and forward crawling resumed (i.e., the AMsh soma activity peak coincided with the end of the reversal behavior, Figs. 2e, g). AMsh activity peaked with a much shorter delay (∼2 s) than in immobilized animals (>60 s). We also measured AMsh soma fluorescence after harsh touch, but there was no correlation between peak fluorescence and any phase of the harsh-touch-induced reversals (Fig. S2). Together, these data support the involvement of AMsh in the nose touch-specific escape behavior, and specifically, that high AMsh activity determines the end of escape reversals.

### Basal AMsh activation is influenced by noxious heat stimuli and is inversely correlated with the duration of harsh touch reversals

We noted that the baseline brightness of the AMsh soma varied by an order of magnitude between individual animals. To test if differences in AMsh soma brightness between animals may have been related to exposure to environmental stressors we compared AMsh baseline fluorescence after incubating animals for 30 minutes under a variety of different established noxious environmental stimuli (e.g., 100mM NaCl, 28°C temperature, starvation, and 4V electric field) to control animals (Thompson *et al*. 2001; Chatzigeorgiou *et al*. 2013; Goodman and Sengupta, 2019). We found no significant difference in AMsh activation for high salinity, starvation, or electric field-treated animals, however, AMsh brightness was significantly lowered in animals exposed to 28°C (Fig. S3c).

We next wondered if AMsh’s basal activity may be predictive of the duration of escape reversals. We compared the AMsh average soma fluorescence during the 30 seconds preceding nose touch with the duration of the resulting reversal. There was no correlation (Fig. 3a), suggesting that nose touch reversal duration is independent of AMsh baseline activity. In contrast, AMsh baseline brightness was inversely correlated with the duration of reversals triggered by anterior harsh touch (Fig. 3b).

**Figure 3:**
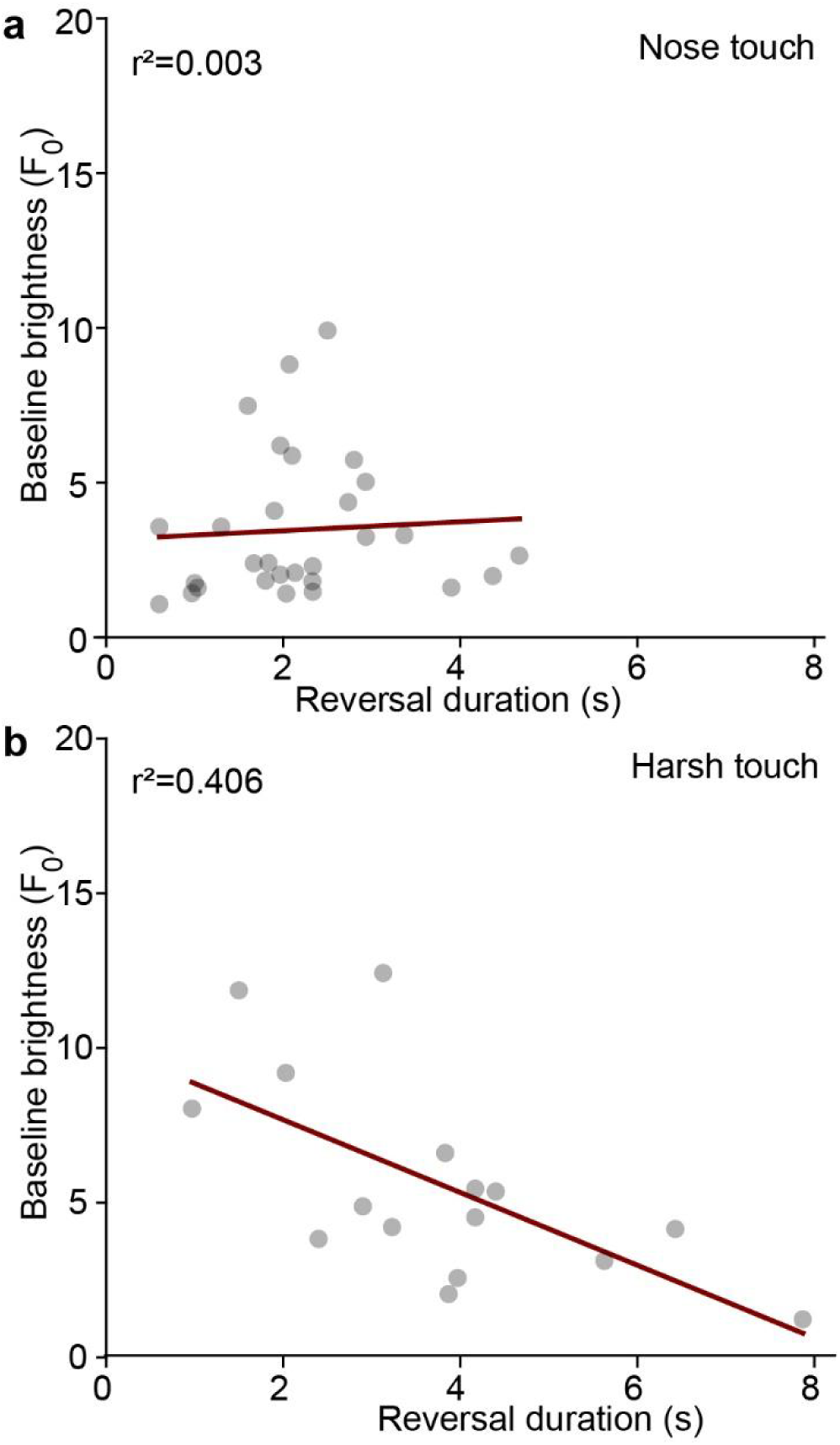
AMsh glia baseline activity is inversely correlated with harsh touch-induced reversal duration. **a)** Average basal AMsh GCaMP6s brightness during 30 seconds preceding a nose touch-induced reversal plotted against the duration of the induced reversal, N= 26. **b)** Average basal AMsh GCaMP6s brightness during 30 seconds preceding a harsh touch-induced reversal plotted against the duration of the induced reversal, N= 17. Spearman correlation indices are shown.

### AMsh calcium signals peak at the end of spontaneous reversals

Our results are consistent with peaks in AMsh soma marking the end of escape reversals specifically initiated by nose touch stimuli. While our initial analysis revealed no correlation between AMsh activity peaks and harsh touch, we wondered whether AMsh activity might be correlated with the end of unprompted (spontaneous) reversals. We thus used freely crawling animals and compared AMsh fluorescence before, during, and after spontaneous (unprompted) reversals. We found that similar to reversals elicited by nose touch, AMsh activity peaked at the end of these spontaneous reversals, but not when reversals started (Fig. 4).

**Figure 4.**
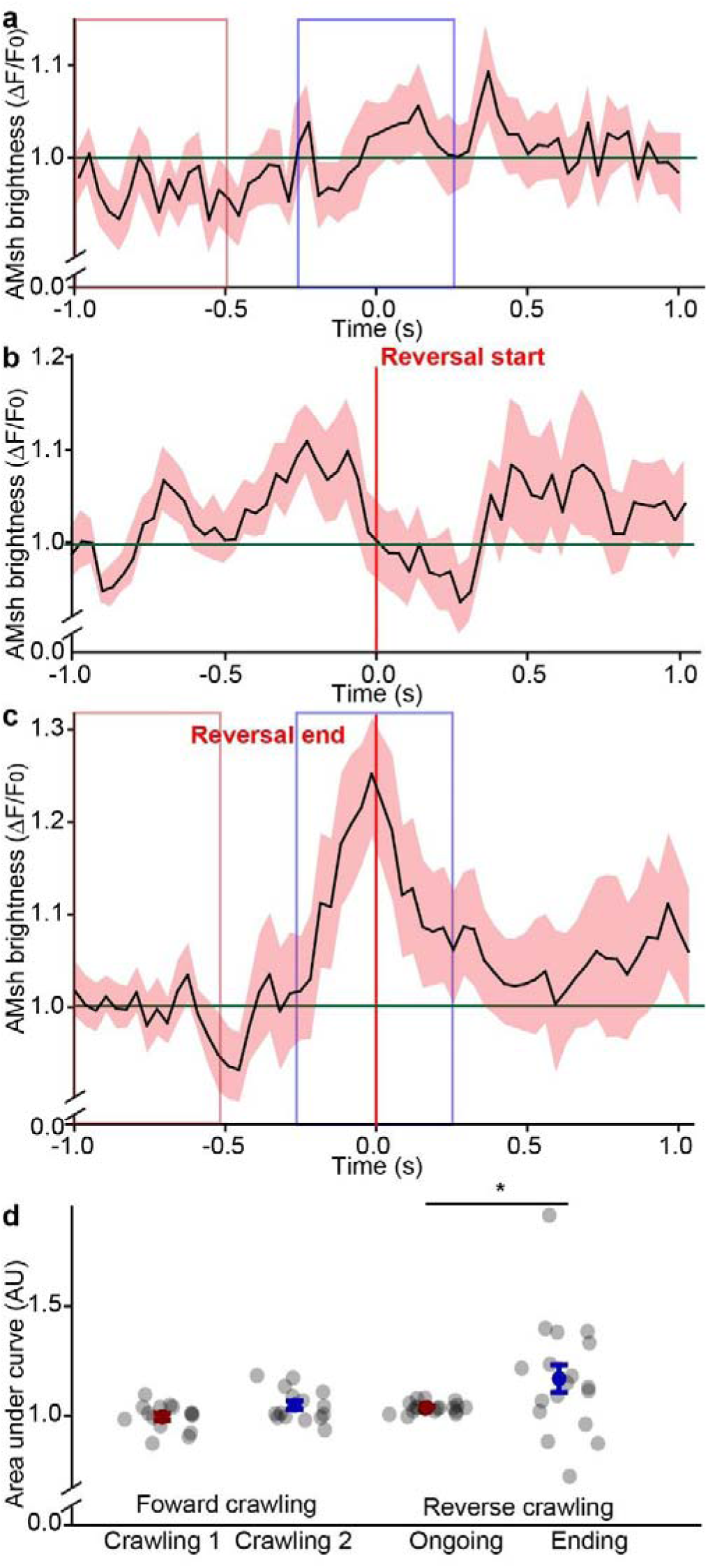
AMsh glia activity peaks at the end of spontaneous (unprompted) reversals. **a)** Average normalized (to background) AMsh GCaMP6s brightness during two seconds of normal forward crawling. For this and the next two plots, the green line indicates the brightness average over the sampled period. **b)** Average AMsh brightness during the second preceding and following the start of a spontaneous reversal. The red line indicates the start of reversal **c)** Average AMsh brightness during the second preceding and following the end of a spontaneous reversal. The red line indicates the end of the reversal. **d)** Comparison of area under the curve for AMsh brightness plots of two 0.5-second intervals during forward crawling bouts in A (indicated by a red and blue shading), and between two 0.5-second intervals during and at the end of spontaneous reversals. Error bars are the standard error of the means (*P< 0.05, N≥10, paired t-tests).

### AMsh nose touch response requires MEC-12

To identify the nose touch transduction mechanisms in AMsh, we searched through the *C. elegans* larva transcriptome library CeNGEN (Taylor *et al*. 2021). We identified several genes involved in mechanotransduction including mechanoreceptors (e.g., *pezo-1*) and other proteins often required for mechano-transduction (e.g. *mec-12*) as being expressed in AMsh glia (Table S1). Because Ding *et al*. (2015) showed that AMsh’s response to nose touch is amiloride-insensitive, we focused our mechanoreceptor search on amiloride-insensitive proteins. We performed a targeted RNA interference (RNAi) screen by targeting the expression of mechanotransduction-relevant genes with expression in AMsh glia as identified by the transcriptome database CeNGEN (Taylor *et al*. 2021). Nose touch-induced AMsh activation in RNAi-treated animals was accomplished on immobilized worms (as in Fig. 1). We applied nose touch stimuli while recording calcium signals in the AMsh terminals (rather than in the more distant AMsh soma) because this is where mechanotransduction of nose touch stimuli is presumed to occur. To evaluate calcium kinetics in the AMsh terminals, we measured peak calcium responses and the area under the calcium decay curve after the nose touch stimulus was applied (see methods).

Reducing expression of the mechanoreceptor-encoding genes identified by CeNGEN to be expressed in AMsh did not alter the nose-touch-evoked calcium responses in these glia (Fig. 5a-d). However, reducing expression of alpha-tubulin (*mec-12*) in AMsh did impair nose-touch responses (Fig. 5). To confirm this finding, we constructed and recorded AMsh nose-touch responses in the AVG30 strain which expresses GCaMP6s in a *mec-12(e1605)* mutant background. Similar to our RNAi results, AMsh response to nose touch response in AMsh was significantly diminished in *mec-12* mutants (Figs. 5e-h).

**Figure 5.**
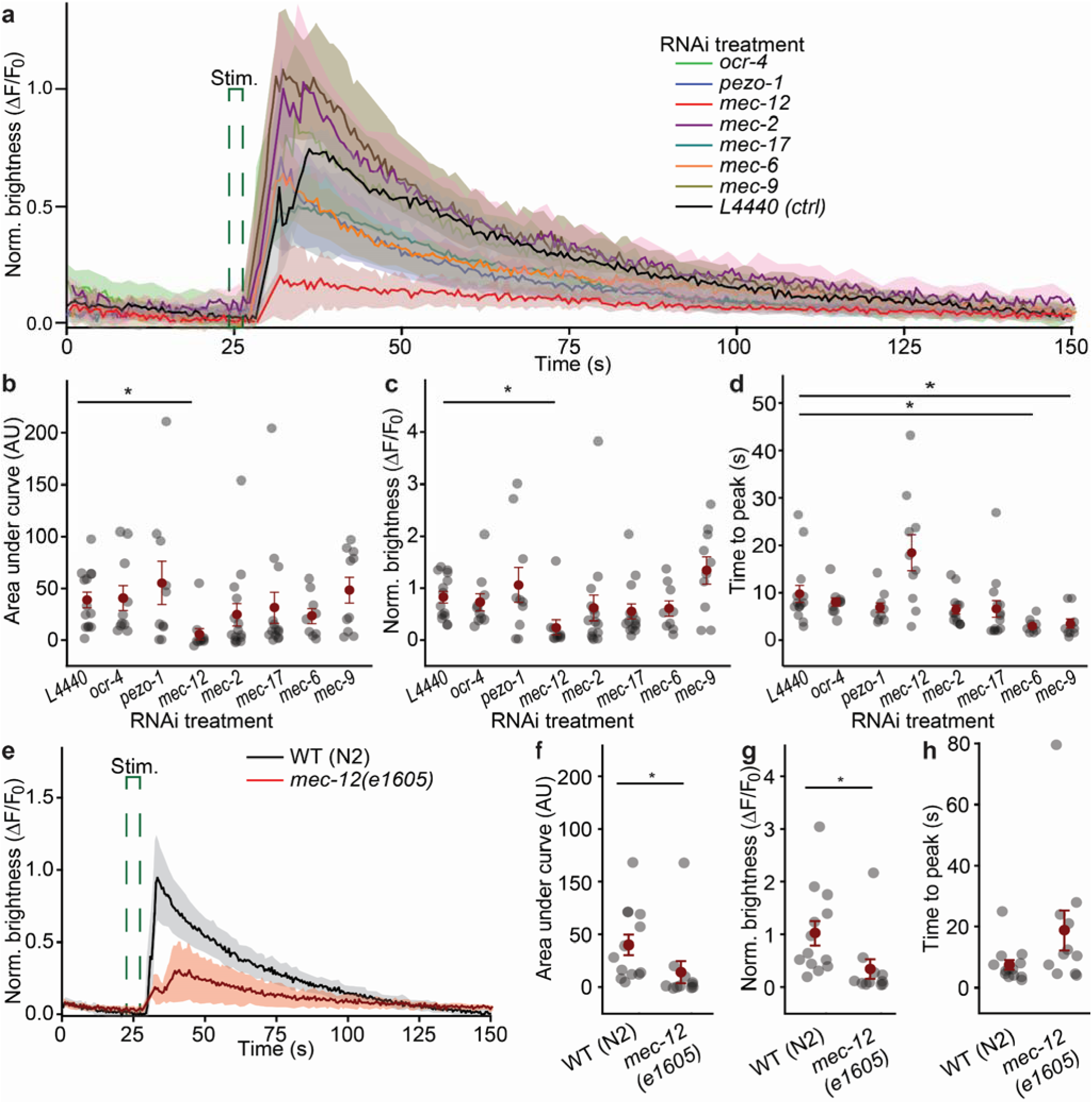
*Mec-12* is required for AMsh glia touch response. **a)** Normalized AMsh terminal brightness (GCaMP6s) response to nose touch stimulation (green dashed line) in glue-immobilized animals following treatment with RNAi targeting genes implicated in touch sensation with known expression in AMsh. **b)** Area under the curve of different RNAi treatments compared to L4440 controls (One-way Anova on ranks, N≥10, H=22.2, *p<0.05, Dunn posthoc test). **c)** Peak AMsh brightness compared to L4440 controls (One-way Anova on ranks, N≥10, H=21.8, *p<0.05, Dunn post-hoc test). **d)** Time to peak brightness compared to L4440 controls (One-way Anova on ranks, N≥10, H=33.5, *p<0.05, Dunn post-hoc test). **e)** Comparison of AMsh brightness between nose-touch wildtype animals (N2) and *mec-12(e1605)* touch-insensitive mutants. **f)** Comparison of areas under the curve for traces in E. **g)** Comparison of AMsh brightness for traces in E. **h)** Comparison of time to peak brightness for traces in E. *p < 0.05, Mann-Whitney rank sum test. Sample sizes are N≥11. For all experiments, data are mean ± SEM.

### Mechanotransduction in AMsh response requires ITR-1 and OSM-9 ion channels

Previous work found that AMsh transduction of noxious odorants required the worm homologs of the mammalian TRPV4 channel (OSM-9) and the inositol triphosphate receptor (ITR-1, Fig. 6a, Duan et al., 2020). Noxious odorants, like nose touch, trigger escape reversals that are mediated by the ASH neurons through a mechanism involving the OSM-9 and ITR-1 (Walker et al. 2009). We again measured calcium influx into the terminals in response to nose touch in worms fed with RNAi bacteria targeting *osm-9* or *itr-1*. We found that RNAi knockdown of either *osm-9* or *itr-1* reduced the amplitude (but not the time to peak) of AMsh calcium responses to nose touch stimuli in immobilized animals (Figs. 6b-e).

**Figure 6.**
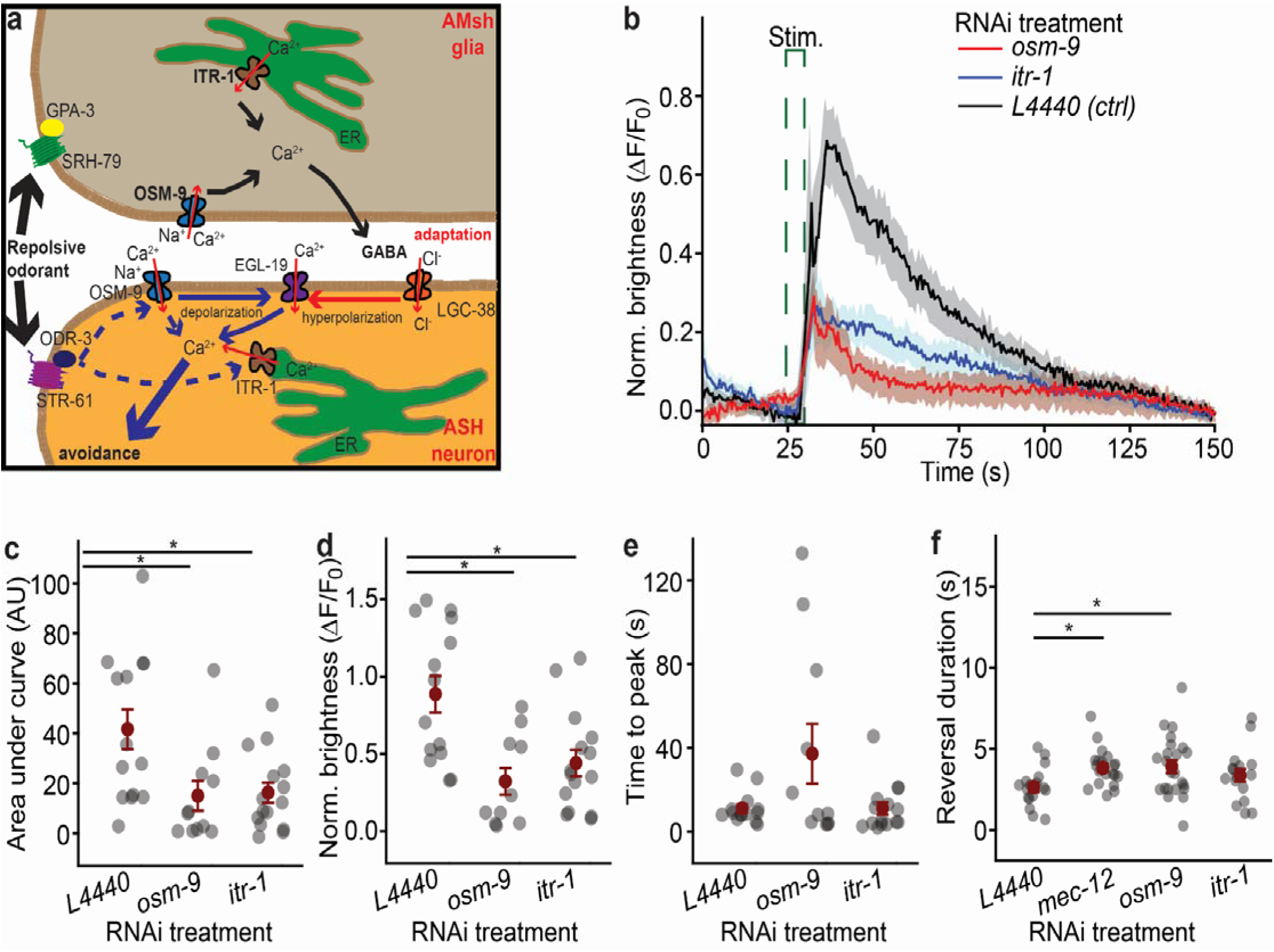
Touch response in AMsh glia relies on *osm-9* and *itr-1* expression. **a)** Working model of AMsh glia-ASH neuron connectivity for an aversive odorant response modeled after Duan *et al*. (2020). **b)** AMsh terminals (GCaMP6s) brightness response to nose touch stimulation (green dashed line) in glue-immobilized animals following treatment with RNAi targeting *osm-9* and *itr-1* expression. **c)** Area under the curve in **b** compared to L4440 controls (One-way Anova on ranks, N≥11, H=9.6, *p<0.05, Dunn posthoc test). **d)** Peak AMsh brightness in **b** compared to L4440 controls (One-way Anova on ranks, N≥11, H=10.7, *p<0.05, Dunn post-hoc test). **e)** Time to peak brightness in **b** compared to L4440 controls (One-way Anova on ranks, N≥11, H=1.2, p>0.05, Dunn post-hoc test). Data are mean ± SEM. *P < 0.05, one-way ANOVA on ranks (N≥11, F_3,66_=5.7, *p < 0.05, Dunn post hoc test). **f)** Duration of nose touch-induced reversals in freely crawling worms, following treatments with RNAi targeting *mec-12, osm-9* and *itr-1* RNAi expression (One-way ANOVA N≥17, F_3,73_=2.7, *p < 0.05, Holm-Sidak post-hoc test). For all experiments, data are shown as mean ± SEM.

To determine if *mec-12*, *osm-9,* and *itr-1* are necessary for normal touch-induced reversal durations in freely moving animals, we used RNAi to reduce their expression. Reduced expression of either *mec-12* or *osm-9* increased reversal durations (L4440 controls = 3.4±1.2 s, *mec-12* = 3.8±1.1 s, *osm-9* = 3.9±1.9 s, One-way ANOVA, F_3,73_= 2.74, Holm-Sidak post hoc, p=0.04, Holm-Sidak post hoc, p=0.03, Fig. 6f). In contrast to nose touch, the duration of reversals triggered by anterior harsh touch was not affected by RNAi targeting of *mec-12*, *osm-9*, and *itr-1* expression (Fig. S4).

### GABAergic signaling is required for normal AMsh modulation of nose touch-induced reversals

Previous studies suggested AMsh glia release □-aminobutyric acid (GABA) to modulate neuronal activity (Duan *et al*. 2020; Fernandez-Abascal *et al*. 2022; Wang *et al*. 2022). To test the potential role of GABA in AMsh modulation of nose touch reversals we used RNAi to reduce the expression of glutamic acid decarboxylase (*unc-25*, required for GABA synthesis). We found that reducing *unc-25* expression non-neuronally (in the N2 background which is refractory to RNAi in neurons but not in AMsh glia) resulted in longer reversals triggered by nose-touch (L4440 controls = 2.4±1.2 s vs *unc-25* RNAi = 3.6±1.3 s, t-test, p<0.001, Fig. 7a). Although all neurons mediating harsh touch-induced reversals express GABA receptors (Gendrel *et al*. 2016), we did not observe a significant increase in reversal duration in response to harsh touch stimulation following RNAi-reduction of *unc-25* expression in this strain (Fig. S5). This suggests that the duration of harsh touch-mediated reversals is determined by different mechanisms than that of nose touch-mediated reversals.

**Figure 7.**
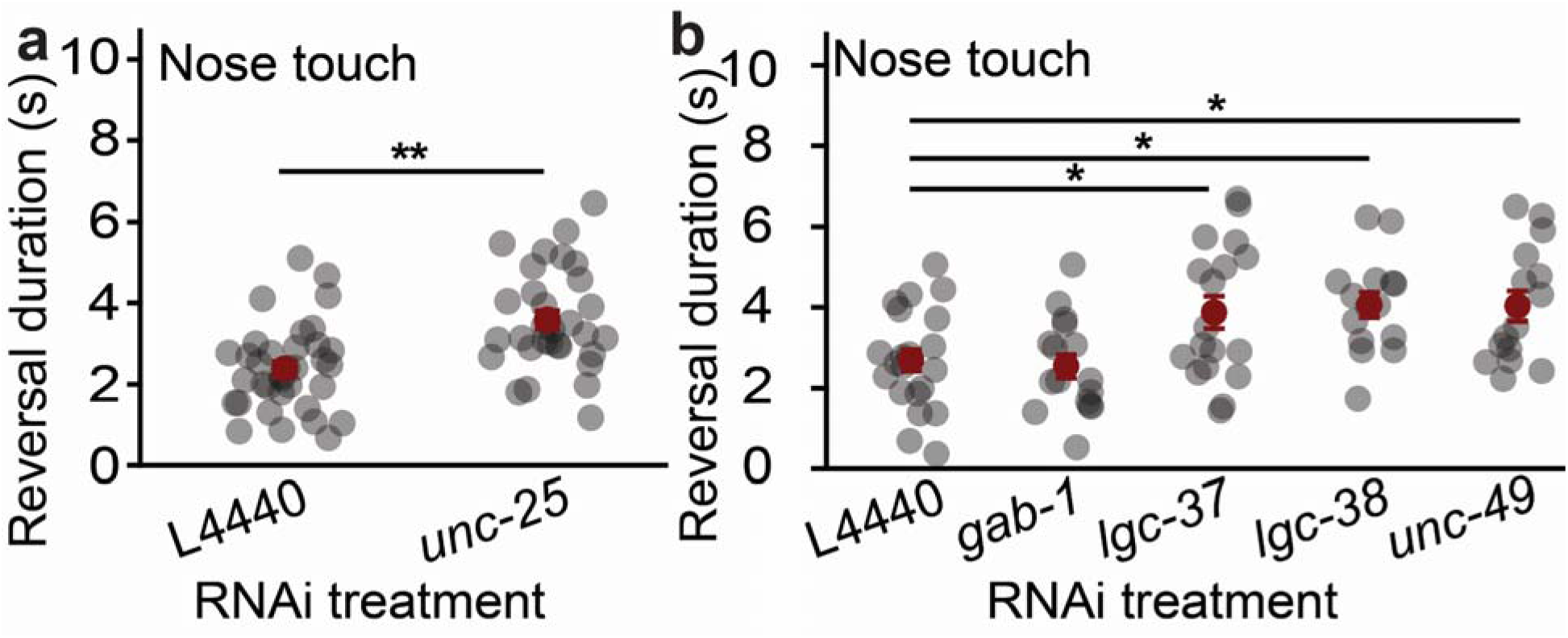
GABAergic signaling is involved in AMsh modulation of touch-induced reversals. **a)** Reversal duration for *unc-25* RNAi-silenced animals in response to nose touch (student’s t-test, L4440 N= 35 and *unc-25* N= 31 **p< 0.001). **b)** Reversal duration for animals with RNAi-silenced GABA receptors in response to nose touch stimuli (one way ANOVA. N≥15, F_4,80_=5.17, *p < 0.05, Holm-Sidak post-hoc test). For all experiments, data are data are shown as mean ± SEM.

All four mechanosensory neurons involved in nose touch responses express GABA receptors. This includes the amphid neurons ASH, but also the OLL, FLP, and CEP which are found outside of the amphids (Gendrel *et al*. 2016, Chatzigeorgiou and Shafer, 2011; Table S2). We used RNAi on a neuronally-RNAi-sensitized strain (TU3311: Calixto et al., 2011) to reduce the expression of GABA receptors (i.e., *gab-1*, *unc-49*, *lgc-37*, and *lgc-38*) known to be expressed in mechanosensory neurons involved in nose touch sensation. Reducing expression of *lgc-37* and *lgc-38* and *unc-49* increased the duration of nose touch-induced reversals (L4440 controls = 2.4±1.3 s, *lgc-37* = 3.9±1.7 s, *lgc-38* = 4.1±1.2 s, *unc-49* = 4.1±1.5 One-way ANOVA, F_4,80_=5.1, Holm-Sidak post hoc, p=0.02, p=0.02 and p=0.02 respectively, Fig. 7b). These data indicate that GABA signaling is involved in mediating AMsh modulation of nose touch-induced reversals.

## Discussion

Previous work implicated AMsh in the modulation of adaptation of ASH to repeated noxious odor (Duang *et al*., 2020) or nose touch stimuli (Ding *et al*. 2015; Chen *et al*. 2022; Fernandez-Abascal *et al*., 2022). While these studies reported AMsh response onset delays on the order of tens of seconds to minutes, none of them addressed the behavioral meaning or potential adaptiveness of such prolonged delayed responses to dangerous stimuli. Our findings corroborate previous observations that AMsh are responsive to nose touch as nose touch-induced calcium waves took tens of seconds to reach the AMsh cell soma (Fig. 1). However, this was only the case in immobilized animals. In contrast, in freely crawling animals, the delay of the calcium wave arrival was dramatically shorter (∼2 seconds), suggesting that the behavioral constraints or continuous sensory stresses animals experience in glue-immobilized conditions alter the functioning of the AMsh glia. We demonstrate that in freely crawling animals the arrival of the calcium peak is associated with a specific component of the escape response, its termination (Fig. 2). This implies that AMsh’s role in the mechanosensation of nose touch stimuli is to modulate the timing of the end of the resulting escape reversal responses. It also fits AMsh’s established role in modulating repeated ASH responses to noxious chemical (Duan *et al*. 2020) and nose touch stimuli (Fernandez-Abascal *et al*., 2022). The modulatory role of the AMsh in mechanosensation appears to be specific to nose touch-induced reversals because no association with anterior harsh touch-induced reversals was found (Fig. S2). Our finding that incubation of animals at noxious temperatures (28°C) was accompanied by a reduction in AMsh brightness (Fig. S3c) further suggests that baseline AMsh activation is affected by at least some external noxious stimuli. However, we did not perform an exhaustive study of environmental factors that could potentially affect AMsh activity. AMsh basal activation had no significant effect on the duration of nose touch-evoked reversals although AMsh baseline was inversely correlated with harsh touch escape duration. It was beyond the scope of this study to establish if this constituted a causal relationship, and if so, through which molecular mechanism the relationship might take place.

Our experiments did not resolve the identity of the mechanoreceptor mediating touch transduction in AMsh. However, we found that the α-tubulin MEC-12 was necessary for normal touch transduction in AMsh (Fig. 5). *Mec-12* is highly expressed in the nervous system of *C. elegans* and in other tissues (including AMsh, Hammarlund *et al*. 2018; Taylor *et al*. 2021). For example, MEC-12 is required for mechanotransduction in all six neurons mediating gentle touch (Fukushige *et al*. 199). It is plausible that MEC-12 plays a similar role in mechanotransduction in the AMsh.

In the ASH neurons, nose touch is primarily mediated by the amiloride-sensitive DEG-1 mechanoreceptor (Geffeney *et al*. 2011). While responses to nose touch in ASH neurons require the TRPV4 channels encoded by *osm-9* and *ocr-2*, they act downstream from DEG-1 in these neurons (Geffeney *et al*. 2011). Mirroring noxious odor transduction, our results show that nose touch responses in AMsh require OSM-9 and ITR-1 (Fig. 6, Duan *et al*. 2020). This is consistent with nose touch sensation being amiloride-insensitive in AMsh (Ding *et al*. 2015) and supports the idea that nose touch in AMsh is mediated by a different mechanosensory mechanism than in ASH neurons. Potential candidates for other TRPV channels mediating mechanotransduction in AMsh are OCR-1 and OCR-2, both known to form functional mechanotransduction channels when associated with OSM-9 (Tobin *et al*. 2002). Alternatively, a different channel might mediate mechanotransduction. If this is the case, OSM-9 would act downstream from that channel like it does in ASH neurons.

RNAi downregulation of GABA synthesis resulted in increased reversal duration in response to nose touch, but not in response to anterior harsh touch (Fig. 7a, S5). Similarly, targeting GABA receptors LGC-37, LCG-38, and UNC-49 (expressed in neurons that mediate nose touch reversals, Table S2) increased the duration of nose touch-induced reversals (Fig. 7b). These data are compatible with a role for AMsh in modulating nose touch reversal duration by releasing GABA to inhibit one or more of these sensory neurons. This is consistent with the proposed modulation by AMsh of ASH responses to repeated noxious odor (Duan *et al*. 2020) and mechanical stimuli Fernandez-Abascal et al. (2022). Further support for AMsh’s use of GABA to modulate neural responses is found through the recently released *C. elegans* transcriptome libraries CenGen and WormSeq, which indicate that AMsh express genes involved in GABA synthesis, transport, vesicle localization to synaptic vesicles, and GABA receptors (see Table S2, Ghaddar *et al*. 2023).

The role of AMsh terminating nose-touch induced reversals appears to rely on the slow progression of distally-generated activation. Nose touch appears to induce activation of the distal processes of AMsh glia and ASH neurons with similar delays (Ding *et al*. 2015). However, the resulting calcium waves propagate to the AMsh soma much slower than in ASH neurons. It, therefore, seems unlikely that AMsh modulation of nose touch sensitive neurons takes place distally in the nose of the animal. Instead, we hypothesize that GABA and Cl^-^ release (Fernandez-Abascal *et al*., 2022) by AMsh and subsequent ASH inhibition takes place closer to the AMsh soma (Fig. s7).

A particular challenge of converging sensory information in polymodal pathways is to retain sensitivity to multiple modalities at once (e.g., how to prevent one modality from decreasing sensitivity to another?). Maintaining high sensory sensitivity is particularly important for pathways underlying escape responses that allow the animal to avoid noxious stimuli. Previous work showed that AMsh’s cell-autonomous responses to noxious odorants are not only slower, but also have a concentration threshold an order of magnitude higher than those of the ASH neurons. AMsh sensitivity to these odors begins at concentrations that already saturate ASH neuronal responses (Duan *et al*. 2020). The hyperpolarization of ASH neurons by AMsh-induced chloride currents could serve the adaptive function of returning these polymodal nociceptive neurons into their responsive range. This adaptation would thus ensure that excitation of ASH neurons by one sensory modality (e.g., a noxious odorant) is not able to impair their ability to respond to noxious stimuli from a different sensory modality (e.g., nose touch), allowing them to continue to trigger escape responses across a broad range of stimuli amplitudes and modalities (e.g. chemical, mechanical, osmotic).

The detection and response to noxious stimuli by polymodal sensory neurons has been shown to be conserved across nematode species (Srinivasan *et al*., 2008). It is possible that glial modulation of polysensory neurons may represent a mechanism to maintain sensitivity in polymodal sensory neurons beyond nematodes. Even though a neuronal connectome for *C. elegans* has existed for decades, it has recently become apparent that extrasynaptic signaling plays a far greater role in neural function than previously believed (Randi *et al*., 2023; Wang *et al*., 2023; and Ripoll-Sánchez *et al*., 2023). It is therefore possible that AMsh modulation of ASH neurons occurs extrasynaptically.

The AMsh of *C. elegans* are remarkable cells that play many important roles in the development and function of the polymodal amphid sensory organ. It is now clear that the role of AMsh glia is not just to serve as support for the dozen sensory neurons in the amphids but also to act as sensors that actively contribute to neural function and the production of adaptive behavior.

## Methods

### Strains

*Caenorhabditis elegans* strains wild-type N2, TU3311, **CB3284** = *mec-12(e1605)* and **OS2248** = *nsls 109 [F16F9.3p::DTA(G53E) + UNC-122::GFP] (genetically ablated AMsh)* were obtained from the Caenorhabditis Genetic Center. The Caenorhabditis Genetics Center (CGC) is supported by the NIH Office of Research Infrastructure Programs (P40 OD010440). AVG26 and AVG30 strains were generated by expressing *F16F9.3p::GCaMP6s* the amphid sheath glia of N2 and CB3284 strains respectively by our lab. All experiments used young day 1 hermaphrodite adult worms reared at 20°C on nematode growth media (NGM) agar plates seeded with *Escherichia coli* strain OP50 strain under standard conditions unless otherwise noted (Brenner, 1974).

### Molecular Biology

We constructed the vector pAVG001 by first replacing the GFP with GCaMP6s in the Ppd9575 plasmid. We next inserted a 2.1-kb promoter of the gene F16F9.3 (FWD 5′-**aagctttatgaaatgcggaacttggagt**-3′ REV 5′-**ggatccctcttactgtcttgggtatttttagagg**-3′, Fung *et al*. 2020) into the pAVG001 vector using HindIII and BamHI restriction enzymes, to make pAVG002 plasmid. RNAi clone targeting *unc-25* and *ocr-4* was generated by restriction cloning about 800 bp and 600 bp coding sequence of *unc-25* (using the primers: using the primers:5’-**ggatccaccggttccatgagaatgtgggaagcagttgg**-3’; 5’-**gggcccggtacccaattgcaggaattcgggattctcaa**-3’) and ocr-4 (with primers: 5’-**ggatccaccggttggatttttagcggatctgg**-3’; 5’-**gggcccggtacccaattgtggatattcgccccaataaa**-3’) respectively in between the AgeI and KpnI cloning sites of L4440 backbone vector as previously described (Nakamura *et al*. 2016), to make pAVG028 and pAVG031 plasmids respectively.

### Transgenic *C. elegan*s strains generated

Germ-line *C. elegans* transformations were performed by standard microinjection methods (Mello *et al*. 1991). We injected the N2 or CB3284 (*mec-12(e1605)*) strain with pF16F9.3::GCaMP6 construct at 50 ng/μL along with 50 ng/μL *Pmyo-3::mCherry::unc-54 3’UTR* as a co-injection marker to generate the AVG26 and AVG30 strains respectively.

### Gene Knockdown

RNA interference by feeding was conducted as previously described (Conte *et al*. 2015). Day 1 adult worms were allowed to lay eggs on RNAi plates for 1 h. The eggs were then grown on the same agar plates containing IPTG and seeded with bacteria containing RNAi empty control vector (L4440), or a vector targeting the gene of interest. Animals were raised to adulthood and tested as gravid Day-1 hermaphrodite adults.

In *C. elegans* neuronal cells are refractory to RNA interference, however, the AMsh are not (Timmons *et al*. 2001; Asikainen *et al*. 2005; Procko *et al*. 2011; Zhang *et al*. 2020) Therefore RNA interference in AMsh was accomplished by using the wild-type (N2) strain, while RNA interference in neurons was accomplished using the pan-neuronally sensitive strain TU3311 (Calixto *et al*. 2010).

All RNAi clones used in this study were obtained from the Fire RNAi library except for *unc-25* and *ocr-4 (*see above). To confirm that RNAi gene knockdown worked as intended, each experiment included a positive RNAi control consisting of N2 worms treated with an RNAi clone targeting *dpy-10* which results in an obvious (dumpy) phenotype involving a shortened body length. This was necessary because silencing gene expression in AMsh glia but not neurons was not predicted to result in a significant decrease in whole-body mRNA levels of the targetted genes.

For the experiments directing gene knockdown to the nervous system (e.g. targeting GABA receptor expression in the TU3311 strain), we additionally used RT-PCR to confirm expression reduction in treatment with a statistically significant impact on nose touch reversal duration (Fig. S6).

### Calcium Imaging

To record AMsh activation we used worms expressing GCaMP6s in AMsh glia in a wildtype background (N2: AVG26) and a *mec-12(e1605)* mutant background (AVG30). Worms were cultivated on standard agar nematode growth media (NGM) plates and tested on day 1 of adulthood.

### Calcium imaging of immobilized animals

AMsh GCaMP6s recordings on immobilized animals was performed on a Zeiss AX10 inverted microscope equipped with an LD Plan-Neofluar 40x/0.6 objective and optiMOS sCMOS camera (Teledyne QImaging, Surrey, Canada). Animals were glued on a glass coverslip coated with 2% NGM agar using WORMGLU (GluStitch Inc., Delta, Canada) immersed in liquid NGM, and mounted on the microscope. GCaMP6s was excited by a Sola Light Engine (Lumencor, Oregon, USA) with a blue light (460-480nm) filter. Touch stimuli averaging 22.46 +/- 4.8 um displacement for 1.2 +/- 0.38 s were delivered to the nose of the worm using a borosilicate glass capillary with a tip diameter of ∼5μm mounted on a Narashige MMO-203 3-axis oil hydraulic micromanipulator (Narashige, Japan).

Each animal was tested once by subjecting them to a single stimulus. Recordings lasted 240 seconds and consisted of a 30-second baseline, a ∼1-sec stimulus, and a 209-second post-stimulus recording. Movies were acquired as a sequence of TIFF images sampled at 2.0 Hz using Micro-manager (Edelstein *et al*. 2010; 2014).

### Calcium imaging of freely crawling animals

Filming of freely crawling worms was performed using a FlyCap2 camera (Point Grey, Vancouver, Canada) mounted on an Olympus SZX12 stereomicroscope, and saved as sequences of TIFF images at 40x magnification and 30Hz sample rate. GCaMP6s was excited using an X-cite 120PC Q light source (Excelitas Technologies, Massachusetts, USA). Two minutes after transfer to filming plate, animals were filmed crawling freely for 2 seconds to establish a baseline, after which a single stimulus was delivered to each animal (either a nose touch or an anterior harsh touch, see section on behavior). Animals were recorded for as long as it took for them to stop reversing and start forward crawling following stimulus administration.

### Image Analysis: immobilized animals

TIFF sequences were uploaded into ImageJ 1.52p (Rasband, 2019). The region of interest tool was used to select areas to be measured (i.e. distal end of AMsh process, mid-process, and AMsh soma, Fig. 1b). The maximum intensity for each area of interest was measured across each TIFF sequence. Areas of interest were monitored and hand-repositioned throughout the stack. The maximum brightness of the background was obtained by dragging the area of interest away from the worm until it lay adjacent to the worm and as close as possible to the measured area in the glia. AMsh brightness was then obtained by subtracting the maximum brightness of the background next to the cell.

AMsh measurements above were normalized by subtracting from them the decay curve of GCaMP6s obtained for each recording using the curve fitting function in SigmaPlot 14.0 (Systat, California, USA). This step converted the recordings from a decay curve to a flat function. To be able to compare calcium kinetics between recordings, we performed one additional normalization operation: We averaged the computed brightness of each cell during the initial 30-second interval (before stimulation) and divided the entire curve by this value as a way to compare all recordings in terms of stimulus-induced changes in brightness over baseline.

During our initial AMsh characterization we recorded from multiple points along the AMsh (Figure 1). However, for later experiments seeking to discern the mechanics of touch transduction, we restricted our measurements to the terminal end of the glia as this region is presumed to be responsible for sensory transduction.

### Image Analysis: freely crawling animals

TIFF sequences were loaded into ImageJ as in the above section. A region of interest encompassing the AMsh soma was hand-selected and measured for each frame of the assay. These selected areas of interest were monitored and repositioned throughout the stack as the animals moved. The maximum brightness of the AMsh soma was obtained along with the maximum brightness of the background, which was obtained by dragging the area of interest away from the soma until it lay adjacent to the worm but as close as possible to the AMsh soma. AMsh soma maximum brightness measurements were normalized by subtracting from them the maximum brightness of the background for each frame of each video.

To compare across recordings, we averaged AMsh brightness of all cells during the initial 1-second interval before stimulation (i.e. 30 frames). We then divided each cell brightness curve by this value so that all recordings had their pre-stimulus brightness normalized to the population mean.

We excluded five frames immediately before and after mechanical stimulation in all our calculations as these contained motion artifacts. Areas under the curve were calculated exempting these frames. To compare between before the end of reversal and during reversal, we used 15 frames during the end of reversal (comprising 7 frames before the end of the reversal, and 7 frames after the end of reversal) and compared it with 14 frames immediately before the end of reversal.

### Behavioral Assays

Freely behaving animals were filmed as a series of TIFF images at a magnification of 40X and a frame rate of 30Hz using a FlyCap2 camera (Point Grey, Vancouver) mounted on an SZX12 stereomicroscope (Olympus) while providing nose touch or harsh touch. We elicited nose touch reversals by placing a platinum wire pick flat on the agar plate, perpendicular to the path of a forward-crawling worm. Forward-crawling animals clashed with this impediment nose-first, causing them to initiate a backward escape. We evoked an anterior harsh touch response by brushing a platinum wire pick against the worm’s top front region of the body. We measured each worm’s reversal length in seconds in response to a stimulus by dividing the number of frames it took the worms to reverse by the sampling rate.

#### Starvation

To assess AMsh calcium signaling under varying satiety conditions, the experimental subjects, AVG26 worms, were transitioned from nutrient-rich NGM plates to unseeded test plates. A 5-minute acclimatization period was allowed, followed by imaging of the worms at 40x magnification using a Point Grey IEEE-1394 camera mounted on an SZX12 stereomicroscope. Starvation conditions were induced by relocating the worms to unseeded NGM plates, where they were left undisturbed for 1 hour before baseline calcium signal assessment. Imaging was conducted in MP4 format and subsequently converted to TIFF format using the Adobe Media Encoder application. TIFF images were then processed in ImageJ, following the methods detailed in the subsequent image analysis subsections.

#### High salinity

To investigate the effect of high salt concentration on AMsh brightness adult AVG26 worms were reared on standard NGM agar plates (i.e. in 50mM NaCl). Baseline measurements of AMsh soma (GCaMP6s) brightness were recorded after transferring animals to an unseeded NGM plate. Animals were allowed to acclimatize for 5 minutes before imaging. Imaging was done at 40x magnification using a Point Grey IEEE-1394 camera mounted on a SZX12 stereomicroscope. Animals were then transferred to NGM agar plates with a salinity of 100mM NaCl, allowed to acclimatize for 5 minutes, and imaged once more as described above. Videos were acquired as MP4 files using a Point Grey IEEE-1394 camera mounted on a SZX12 stereomicroscope at 40x magnification. These MP4 videos were converted to TIFF images using the Adobe Media Encoder application and AMsh soma brightness was manually measured and normalized to background brightness using ImageJ (FIJI), following the methodologies detailed in the image analysis subsections.

#### Heat-Shock

To investigate AMsh response to change in temperature, 10 randomly chosen adult AVG26 worms were grown under standard room temperature (20°C). The brightness of the AMsh soma (GCaMP6s) was recorded for each worm while crawling freely on agar plates seeded with OP-50. After recording the pre-heat-shock measurements, the worms were transferred to newly seeded agar plates. To ensure individual identification and alignment of pre-and post-treatment measurements, a copper ring was positioned around each worm on the new plates. The worms within the copper rings were placed in an incubator at 28°C for 30 minutes. After 30 minutes, the plates were removed from the incubator and the brightness from the AMsh soma were recorded immediately for each worm. All Imaging was done at 40x magnification using a Point Grey IEEE-1394 camera mounted on a SZX12 stereomicroscope. Videos were acquired as MP4 files using a Point Grey IEEE-1394 camera mounted on a SZX12 stereomicroscope at 40x magnification. These MP4 videos were converted to TIFF images using the Adobe Media Encoder application and AMsh soma brightness was manually measured and normalized as described above.

#### Electric fields

Electric fields were generated by generating a voltage between two probes inserted at opposing ends of an NGM agar plate (which contains 50mM NaCl) as previously described (Chrisman *et al*. 2016). A constant electric field strength of 4V was delivered through metal probes powered by a LONG WEI DC power supply (model PS-305D). Baseline measurements of AMsh soma (GCaMP6s) brightness were recorded before and during the exposure to electric fields. Videos were converted to TIFF images and AMsh soma brightness was measured as described above.

### Statistical analysis

All statistical analyses were performed using Sigmaplot 14.5 software from Systat Software, Inc. Paired comparisons were assessed through paired t-tests. In contrast, comparisons between two distinct treatments were examined using student’s t-tests under the conditions of normal data distribution and similar variances. When normality and equal variance tests failed, comparisons were conducted via the Mann–Whitney Rank Sum Test. Multiple parametric datasets were compared using One-Way ANOVA, followed by Holm–Sidak all-pairwise post hoc tests. For non-parametric datasets, Kruskal-Wallis One-Way ANOVA on Ranks was employed, followed by Dunn’s post hoc tests, accounting for different sample sizes within the groups. The standard convention for significance was adopted (i.e., *P < 0.05 and **P < 0.001). Power analysis was carried out using Sigmaplot 14.5 for parametric tests, and StudySize 3.0 by CreoStat HB (Sweden) for non-parametric tests, where the highest standard deviation and the smallest sample size were utilized for minimum detectable mean differences when groups had unequal sample sizes. Please refer to Supplementary Spreadsheet 01 and 02 for raw data and statistics for each figure in this study (respectively). Raw video recordings obtained for this study will be available through the server Zenodo at: 10.5281/zenodo.10372774.

---

**Figure S1:**
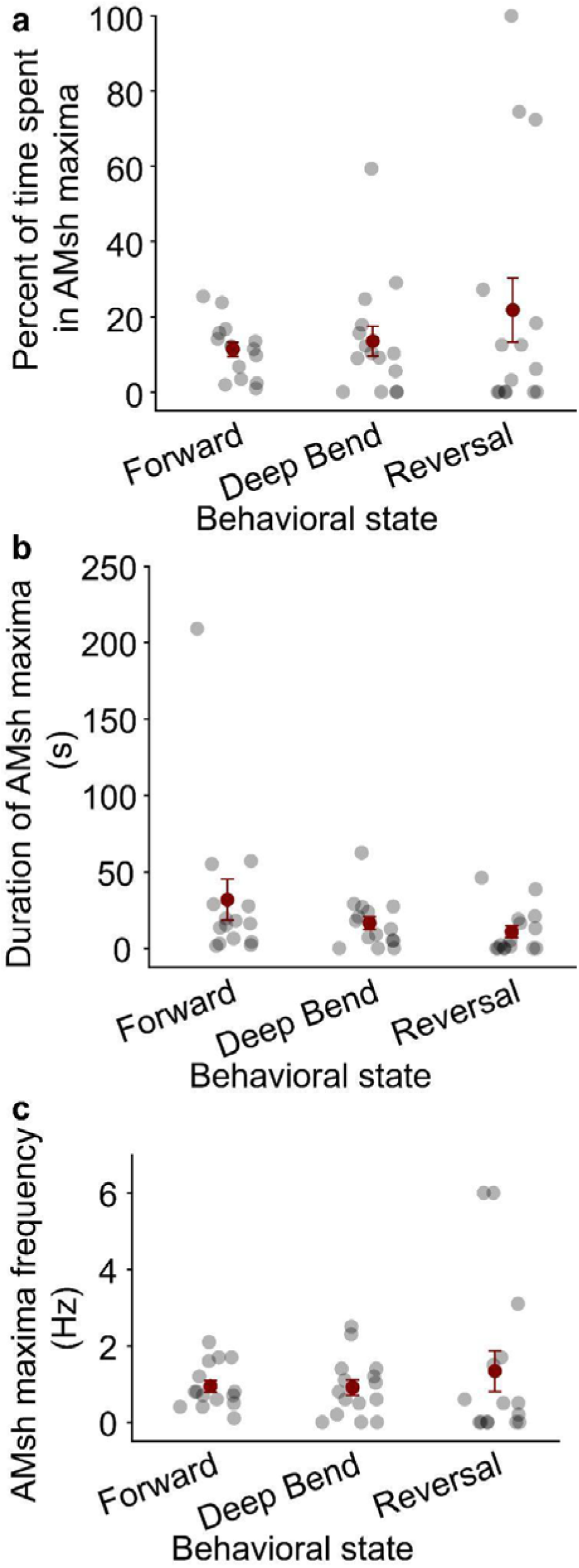
Peaks in AMsh soma brightness are not correlated with crawling or turning behaviors. AMsh soma brightness maxima are defined as GCaMP6s brightness exceeding its average brightness by more than 1 StDev in freely crawling worms (see methods). **A)** There was no difference in the percent of the time occupied by AMsh maxima between different locomotor behaviors (One Way ANOVA on ranks, N=15, H_2_=0.36, P= 0.8). Similarly, we did not measure a significant difference between the duration (One Way ANOVA on ranks, N=15, H_2_=4.38, P= 0.1) **(B)** or frequency (One Way ANOVA on ranks, N=15, H_2_=1.39, P= 0.5) **(C)** of AMsh peaks during the performance of these behaviors.

**Figure S2.**
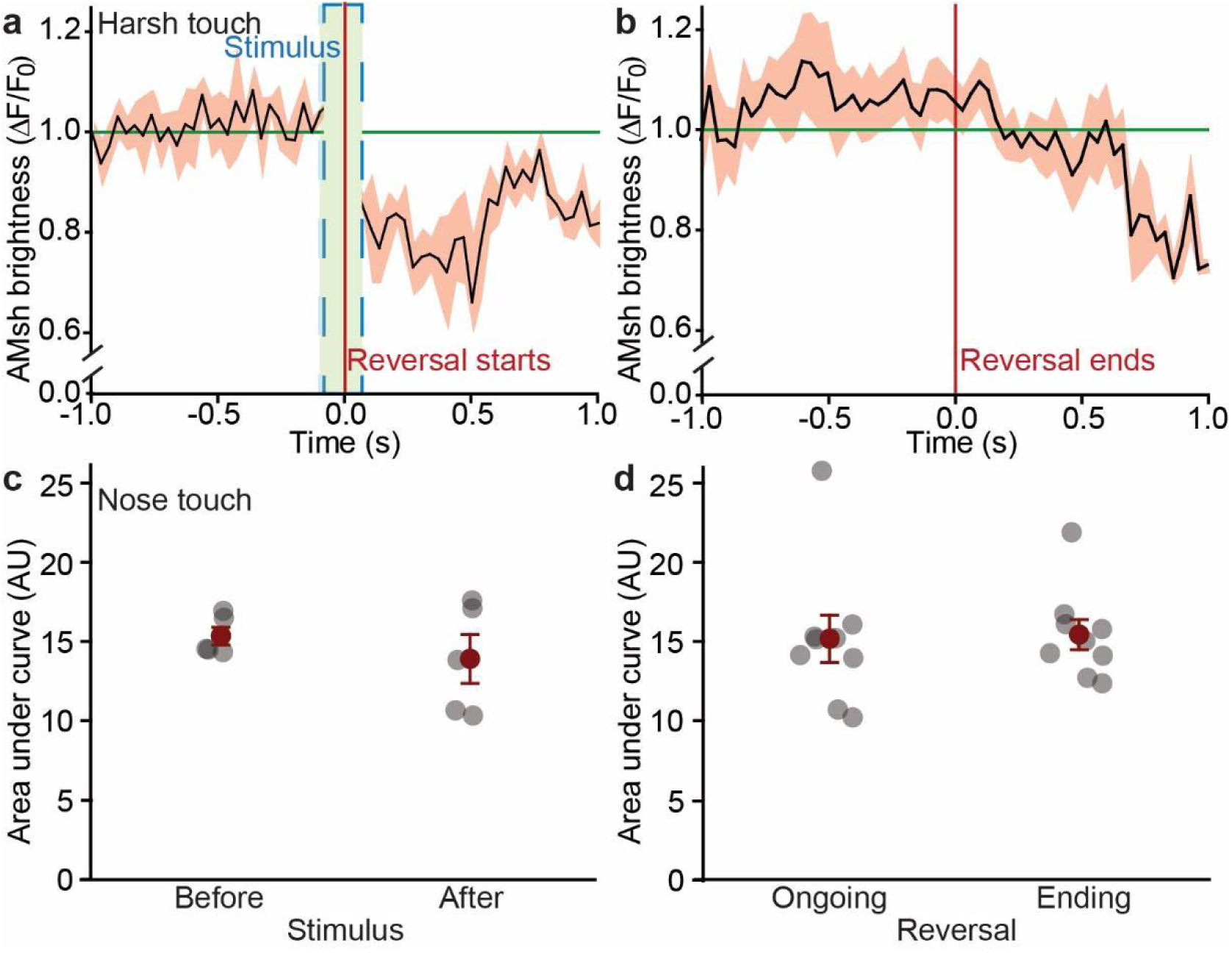
AMsh glia activity does not peak at the end of harsh touch-induced reversal **a)** Normalized (to background) maximal GCaMP6s brightness in the AMsh glia of freely crawling worms in the second before and after administration of anterior harsh touch (green dashed lines). The red vertical line demarcates the time when the worms initiated their characteristic escape reversal. **b)** AMsh brightness in the second before and after the end of the nose touch-induced reversals. The vertical red line denotes the time when worms ceased backward motion and resumed forward motion. Comparison of the area under the curve for AMsh GCaMP6s brightness for the 30 milliseconds before and after anterior **(c)**, and before and during the end of the elicited backward locomotion for nose touch **(d)**. Data are shown as mean ± SEM. *P < 0.05, paired student’s t-tests. N=11.

**Figure S3:**
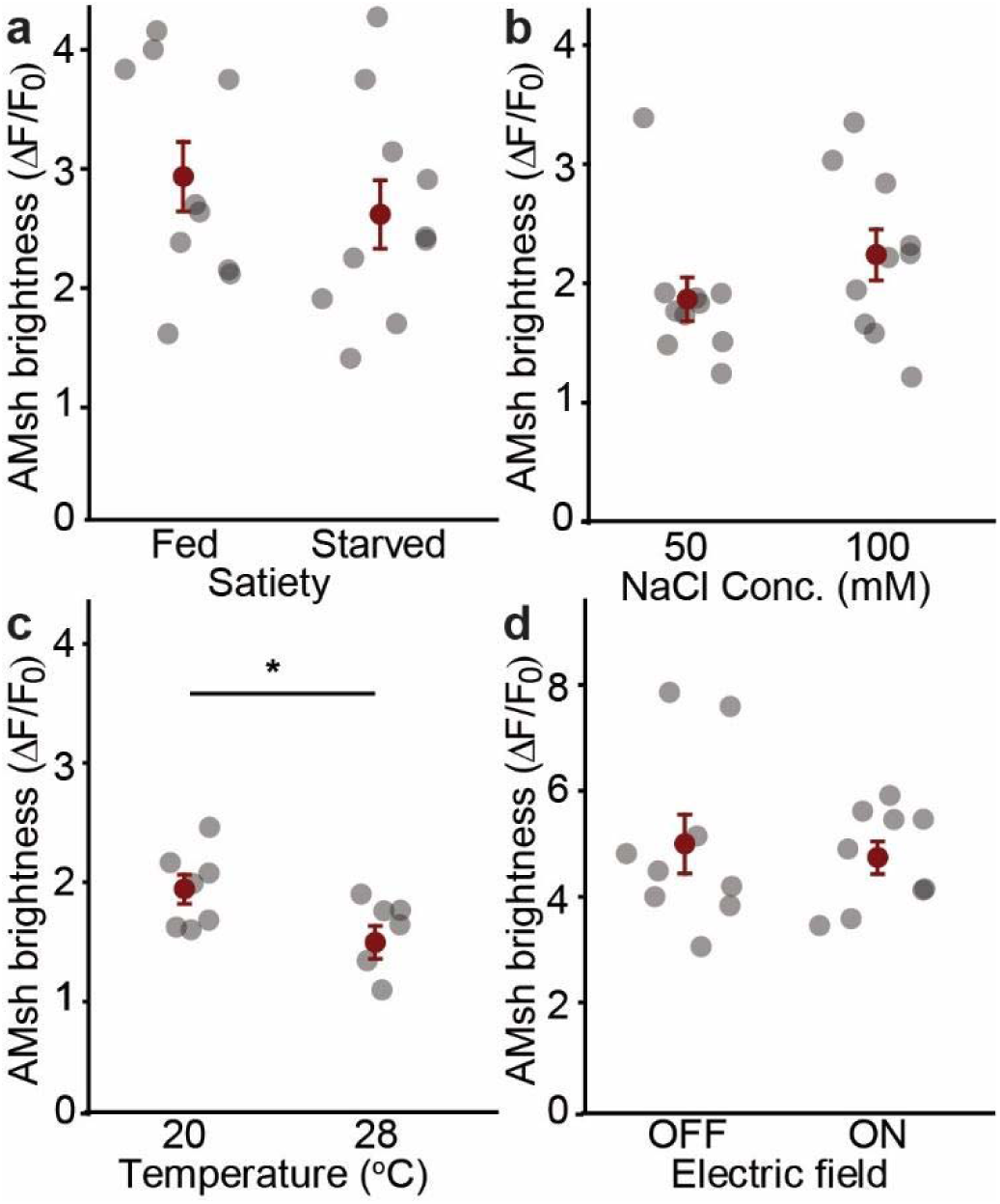
AMsh glia GCaMP6s baseline activity in different stress conditions. **a)** Before and after starvation for 30 minutes. N=10. **b)** Before and during exposure to higher NaCl concentration (100mM). N=10 **c)** Before and after being heated to 28°C. N=7. **d)** Before and during exposure to electric fields. N=10. For all experiments, data are shown as mean ± SEM. *P < 0.05, paired student’s t-tests.

**Figure S4.**
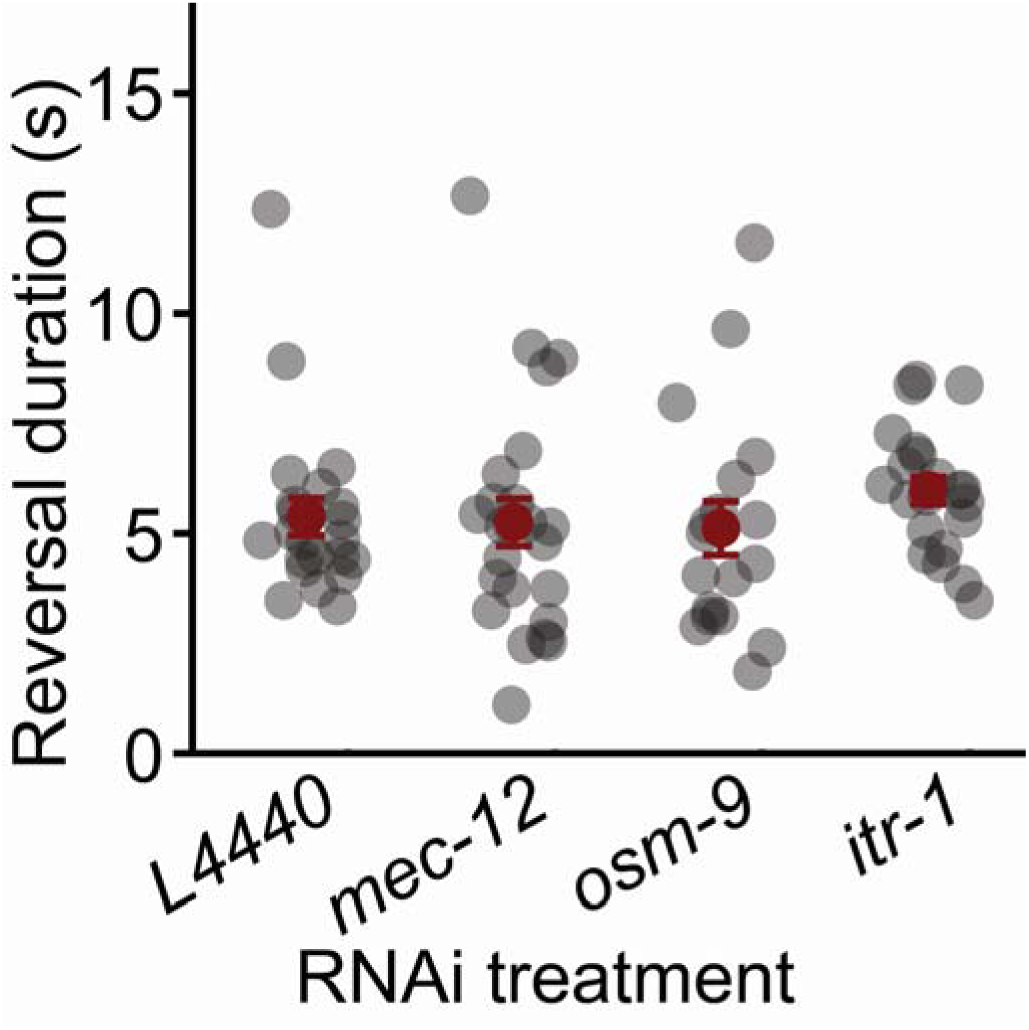
Duration of anterior harsh touch-induced reversals following treatments with RNAi targeting *mec-12, osm-9* and *itr-1* RNAi expression, in comparison to control (one way ANOVA on ranks. N≥18, H_3_=6.11, P = 0.11).

**Figure S5.**
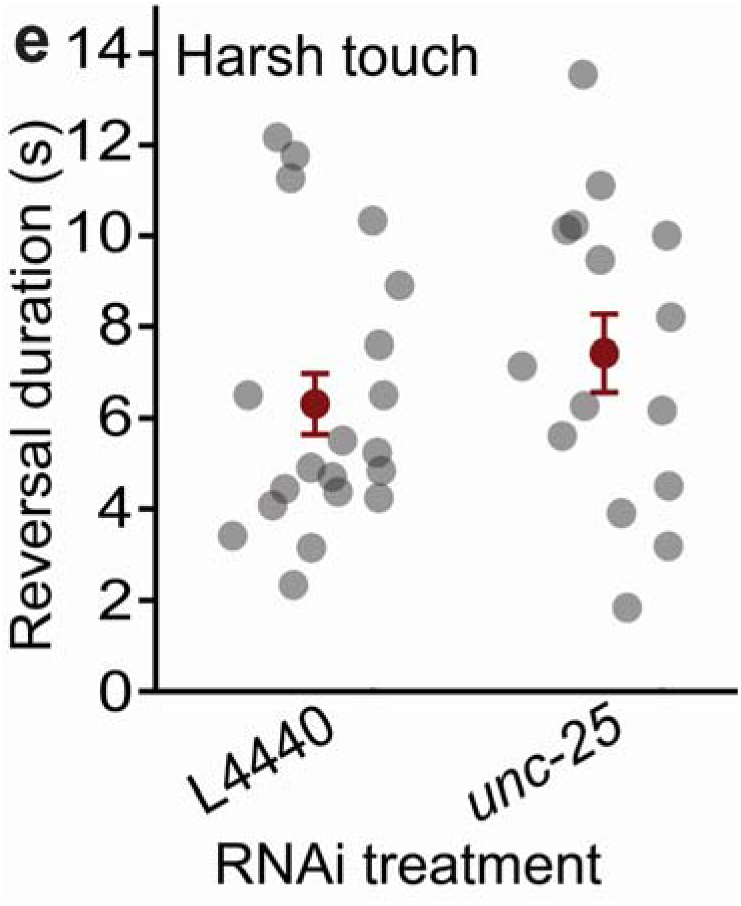
Silencing *unc-25* does not prolong harsh touch-induced reversal. (Student’s t-tests, N≥15, P=0.23). Data shown as mean ± SEM. *P < 0.05, paired student’s t-tests.

**Figure S6.**
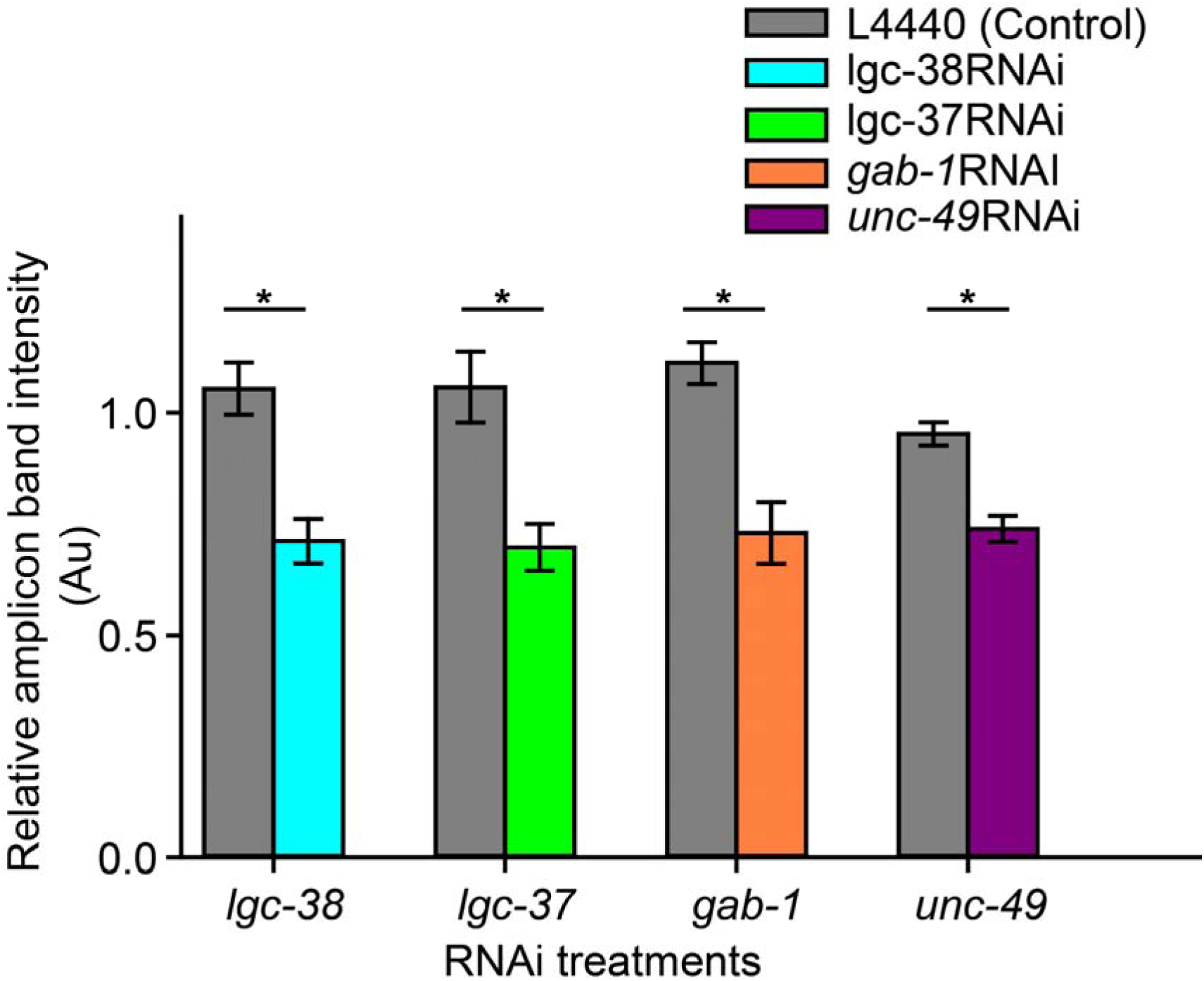
Quantification of the RT-PCR products for GABA receptor genes *lgc-38*, *lgc-37*, *gab-1* and *unc-49* in empty l4440 (grey bar) vector control and RNAi treated TU3311 worm strain relative to *pmp-3* housekeeping gene (Student’s t-tests, N=6, *P<0.05). Data shown as mean ± SEM.

**Figure S7.**
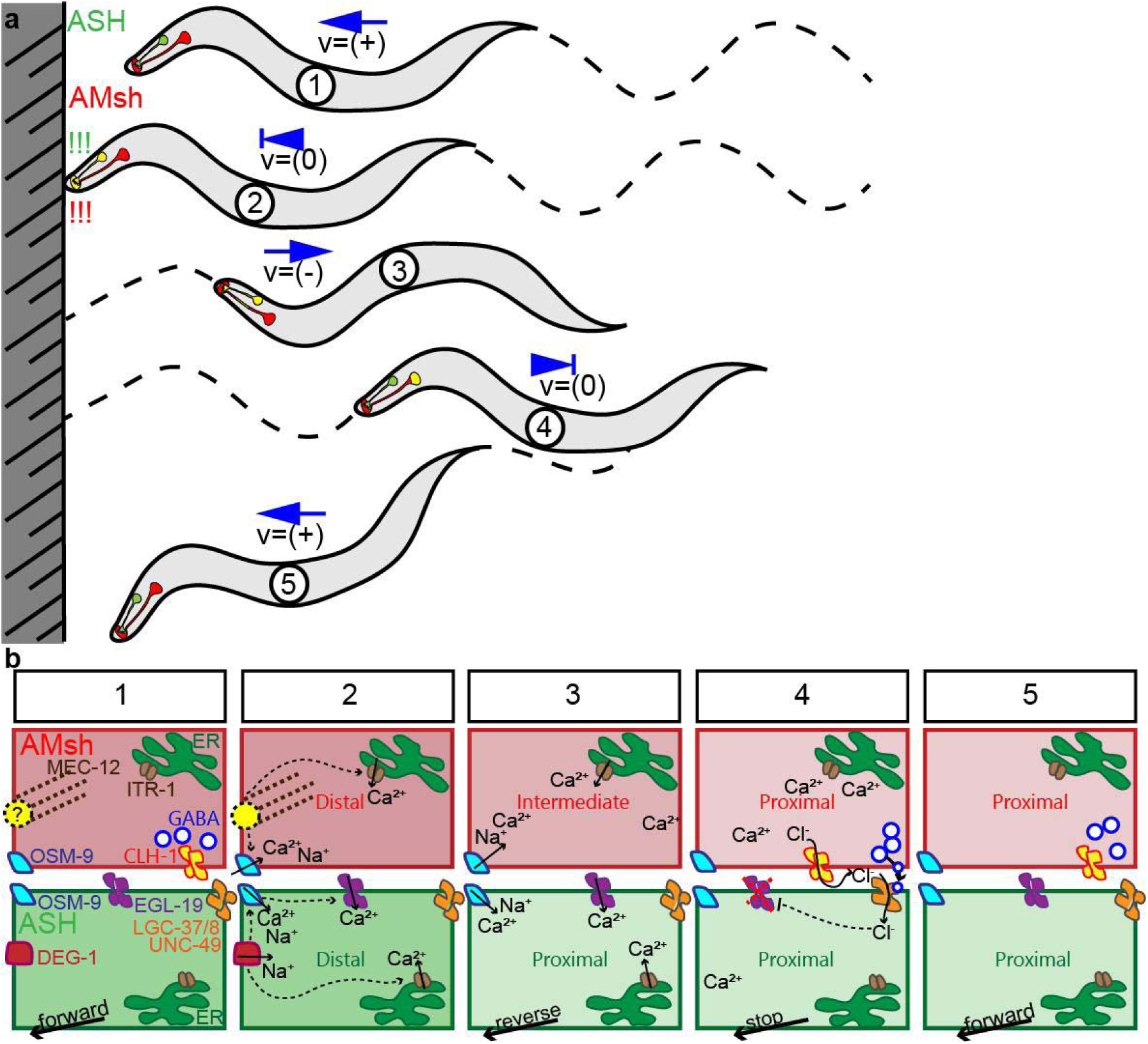
Behavioral and molecular models for glial modulation of nose touch reversal duration. **a)** Based on our results and previous work by Fernandez-Abascal et al. (2022), and Ding et al. (2015) we propose a model where a forward crawling worm colliding with an immobile object (1) simultaneously activates ASH neurons and AMsh glia (2). (3) ASH activation propagates from periphery to soma faster than in AMsh glia resulting in the initiation of a backwards crawling escape response. (4) Sometime after the initiation of the reversal escape, AMsh activation reaches its soma causing AMsh to inhibit ASH and terminate the ongoing reversal. (5) Once the escape response is terminated forward crawling can resume. **b)** Proposed molecular mechanism responsible for the behavior described in a. 1) Nose touch activates ASH neurons through DEG-1 mechanoreceptors and AMsh glia through (unidentified) receptors (involving alpha tubulin MEC-12). 2) At the ASH neurons terminals, mechanoreceptor activation causes influx of extracellular calcium through OSM-9 and EGL-19 channels, and of endoplasmic reticulum calcium through ITR-1 channels. This results in the termination of forward locomotion and initiation of backwards crawling. Simultaneously, mechanical activation of mechanoreceptors on AMsh glia cause activation of OSM-9 and ITR-1, resulting in the increased calcium at the AMsh terminals. 3) The distal increase in cytosolic calcium at the AMsh terminals slowly travels towards the soma (through OSM-9) and ER calcium (through ITR-1). 4) Once the calcium wave in AMsh arrives at their soma, it causes the release of GABA from AMsh glia which binds and activates LGC-37/38/UNC-49 (GABAAR) chloride channels in ASH which then open to allow influx of chloride released by AMsh through CLH-1 channels. The influx of chloride into ASH neurons hyperpolarizes them, closing their voltage gated (EGL-19) calcium channels. ASH hyperpolarization terminates backwards locomotion. 5) With ASH neurons hyperpolarized forward locomotion resumes.

**Table S1:** List of mechanoreceptors and other genes required for mechano-transduction that are expressed in AMsh according to the CeNGEN (Taylor *et al*. 2021).

| Gene name | Expression level in AMsh<br>(Source: CeNGEN) | Family | Notes |
| --- | --- | --- | --- |
| *mec-12* | 926.710 | Alpha-tubulin | accessory proteins |
| *mec-17* | 518.254 | Alpha-tubulin | accessory proteins |
| *mec-9* | 58.53 | Uncharacterized protein | accessory proteins |
| *unc-8* | 110.035 | Deg/ENaC | amiloride sensitive |
| *deg-3* | 16.801 | Deg/ENaC | amiloride sensitive |
| *egas-4* | 9.137 | Deg/ENaC | amiloride sensitive |
| *mec-2* | 9.137 | Stomatin | accessory proteins |
| *ocr-4* | 5.961 | TRPV | amiloride insensitive |
| *mec-6* | 1.987 | Paraoxonase | accessory proteins |
| *pezo-1* | 1.987 | Piezo | amiloride insensitive |

**Table S2:** List of GABAergic genes expressed in AMsh glia and nose touch sensitive neurons according to CeNGEN and WormSeq transcriptomic databases (Taylor *et al*. 2021, Ghaddar *et al*. 2023).

| Cell | Gene | Expression level<br>CeNGEN (larvae) | Expression level<br>WormSeq (adult) | Notes |
| --- | --- | --- | --- | --- |
| AMsh | *unc-25* | 599.811 | 42.29 | glutamate decarboxylase |
| AMsh | *gta-1* | 195.717 | 108.17 | 4-aminobutyrate aminotransferase |
| AMsh | *unc-46* | 0 | 12.89 | Localizes UNC-47 to vesicles |
| AMsh | *unc-47* | 0 | 27.09 | Vesicular GABA transporter |
| AMsh | *unc-30* | 0 | 1.01 | *unc-25* regulator |
| AMsh | *snf-11* | 0 | 16.48 | Plasmallema GABA transporter |
| AMsh | *lgc-37* | 0 | 0.45 | GABA A receptor |
| AMsh | *lgc-38* | 73.795 | 1.33 | GABA A receptor |
| AMsh | *unc-49* | 4.278 | 9.92 | GABA A receptor |
| AMsh | *exp-1* | 0.195 | 12.41 | Excitatory GABA receptor |
| ASH | *gab-1* | 0.993 | 4.55 | GABA A receptor |
| ASH | *lgc-37* | 1.73 | 0 | GABA A receptor |
| ASH | *lgc-38* | 4.746 | 0 | GABA A receptor |
| ASH | *unc-49* | 10.878 | 4.16 | GABA A receptor |
| CEP | *exp-1* | 0.545 | 35.84 | Excitatory GABA receptor |
| CEP | *gab-1* | 12.715 | 0 | GABA A receptor |
| CEP | *lgc-38* | 281.285 | 0 | GABA A receptor |
| CEP | *unc-49* | 15.493 | 77.51 | GABA A receptor |
| FLP | *gab-1* | 21.368 | n/a | GABA A receptor |
| FLP | *lgc-37* | 15.354 | n/a | GABA A receptor |
| FLP | *lgc-38* | 156.986 | n/a | GABA A receptor |
| OLQ | *lgc-38* | 1128.494 | 648.38 | GABA A receptor |
| OLQ | *unc-49* | 106.508 | 0 | GABA A receptor |

